# Identification, antimicrobial resistance and molecular detection of Shiga toxin-producing *E. coli* (STEC) isolated from raw milk of cattle in Chitwan, Nepal

**DOI:** 10.1101/2024.01.30.578021

**Authors:** Sunita Shrestha, Rebanta Kumar Bhattarai, Hom Bahadur Basnet, Himal Luitel, Sirjan Bastola

**Affiliations:** Department of Veterinary Microbiology and Parasitology, Agriculture and Forestry University, Rampur, Nepal; Center for Biotechnology, Agriculture and Forestry University, Rampur, Nepal; Department of Veterinary Microbiology and Parasitology, IAAS, Paklihawa Campus, Tribhuvan University, Paklihawa, Nepal

**Keywords:** Shiga toxin-producing *Escherichia coli* (STEC), O157:H7 STEC, non-O157:H7 STEC, *stx* gene, raw cattle milk, antibiogram

## Abstract

Shiga toxin-producing *Escherichia coli* (STEC) is one of the most common foodborne pathogens leading to gastroenteritis with symptoms of mild diarrhoea to the potential life-threatening haemolytic uremic syndrome. Serotype O157:H7 has been the most predominant type worldwide but there are an increasing number of outbreaks caused by non-O157:H7 STEC serotypes. This is the first study about STEC in milk used in Nepalese communities. A total of 267 raw milk samples were taken from different sites in Chitwan District, one of the major milk-producing districts of Nepal, using multi-stage sampling. The samples underwent microbiological and molecular analysis. The overall prevalence of *E. coli* was found to be 29.6% (79/267) in raw cattle milk. *Z3276* gene, a specific genetic marker of *E. coli* O157:H7 was detected only in three isolates which also carried *stx* genes. The overall prevalence of STEC in raw cattle milk was 11.6% (31/267) whereas the prevalence of STEC among *E. coli* isolates was 39.2% (31/79). The prevalence of O157:H7 STEC is 1.1% (3/267) and of non-O157:H7 STEC is 10.5% (28/267). Out of 12 antibiotics tested, the sensitivity of STEC to an antibiotic was highest with azithromycin, and levofloxacin, and the most resistant was with tetracycline, and doxycycline. The findings revealed that raw cattle milk was contaminated with STEC, posing a potential public health concern. The discovery of a greater occurrence of *stx2* genes in this study highlights the possibility of more severe complications in humans. Molecular-based STEC surveillance is essential to minimize the risk of its transmission.

## INTRODUCTION

Shiga toxin-producing *Escherichia coli* (STEC) stands as one of the most prevalent food-borne pathogens (1) causing gastroenteritis (2). Microbial food-borne illness still remains a global concern despite the extensive scientific progress and technological developments achieved in recent years in developed countries (3). Food-borne diseases occur commonly in developing countries because of the prevailing poor food handling and sanitation practices, weak regulatory system, inadequate food safety laws, lack of education for food handlers and lack of financial resources to invest in safer equipment (4).

STEC is considered as a worldwide threat to public health (5, 6) and it has several virulence factors such as Shiga toxins (Stx1 and Stx2), intimin (*eaeA*) and enterohemolysin (*hlyA*). Among diarrhoeagenic *E. coli*, STEC includes the most virulent strains (7). STEC has genes encoding for the production of Shiga toxins (Stx) in their genomes (8), a toxin very similar to that of *Shigella dysenteriae* (9). STEC is a zoonotic pathogen that is a major cause of diarrhoea worldwide (10). STECs are pathogens capable of causing sporadic and epidemic infections in humans. Symptoms of STEC infections range from mild diarrhoea to life-threatening hemolytic uremic syndrome (11). The first recorded case of food-borne illness associated with *E. coli* O157:H7 was an outbreak of haemorrhagic colitis from ground beef patties in 1982 (12). Serotype O157:H7 STEC has been the predominant type worldwide (13). However, there has been a notable increase in the number of outbreaks caused by non-O157 STEC serotypes (14). *Z3276* was identified as a specific marker of *E. coli* O157:H7 (15, 16).

Altogether more than 200 STEC serotypes have been reported and more than 100 have been linked with human infection (17). There have been several very large outbreaks around the world and their public impact has often been pronounced. Every year there have been minimum of two STEC outbreaks since 2006 to till date (18) Ruminants, with cattle being the primary reservoir, are considered important carriers of STEC (19-22). Sporadic cases and outbreaks of human diseases caused by food-borne pathogens and STEC have been linked to vegetables, unpasteurized fruit juices, water, foods of animal origin, such as meat, milk, and dairy products (23-25). Contaminated raw milk is one of the primary sources of food-borne illnesses (26). Raw milk, being a nutritious food, serves as an ideal medium for the growth of various bacteria including pathogenic organisms which have a significant impact on public health (27). Many microorganisms can get access to milk and milk byproducts among which *E. coli* is recognized to be of primary concern (28). Among 81 raw milk outbreaks during 2007-2012, STEC was the causative agent of 17% of outbreaks, making it the second most common pathogen following *Campylobacter* spp. (29). The presumptive route of STEC in milk is through faecal contamination and also direct shedding from an infected udder (30).

Different antibiotic resistance profiles have been detected in *E. coli* isolates from different sources, including humans, animals, and foods (31). Unpasteurized milk is considered a main source of *E. coli* isolates resistance to multiple antibiotics (32). *E. coli* can become resistant due to the habit of excessive use of antibiotics (33). One of the most controversial areas in the management of STEC infections lies in the possible effect of antimicrobials on the natural history of the infections. As no specific drug has demonstrated effectiveness, the treatment of STEC is mainly symptomatic (34, 35). Because antimicrobials may lyse bacterial cell walls, thereby liberating Shiga toxins (36), and/or cause increased expression of Shiga toxin genes in vivo (37), they are not recommended for treating STEC O157:H7 infections (38). The antibiotics do not benefit the infected but they are associated with an increased likelihood of developing HUS so prevention efforts should prioritize primary infection, prevention, and managing extra-intestinal consequences (39). Although STEC infections are not aggressively treated with antimicrobial therapy and many isolates are susceptible to numerous antimicrobials, some reports also indicate that antimicrobial resistance of STEC is on the rise (40). So, we aim to find the antibiogram of *E. coli* and STEC in Chitwan, Nepal.

Nepal is an agricultural country and cattle farming is one of the main sources of income. Chitwan, one of Nepal’s top five districts for the highest total milk production, (41) has now become fully sufficient for milk production (42). Previous studies have revealed that subclinical mastitis is prevalent in Nepal, with *E. coli* as a commonly associated pathogen (43, 44). However, there have been no studies to determine the prevalence of Shiga toxin-producing genes in *E. coli* in Nepal. Subclinical mastitis is a condition in animals that typically lacks noticeable symptoms and often remains undetected. Consequently, milk affected by subclinical mastitis can be consumed by consumers without their awareness of the issue. Cattle and raw milk are considered primary sources of STEC and since *E. coli* is known as the most commonly associated organism with subclinical mastitis, it is crucial to investigate the prevalence of Shiga toxins in raw milk. So, the aim of this study is to provide insights into the prevalence and antibiogram of Shiga toxin-producing *E. coli* (STEC) from raw cattle milk in Nepal.

## RESULTS

Out of 267 milk samples tested, biochemical tests determined 74 isolates as *E. coli* and five isolates as *E. coli* O157:H7 which results in the overall prevalence of *E. coli* to be 29.6% (79/267) in raw cattle milk from Chitwan.

### Detection of *Z3276* gene

Among five isolates of *E. coli* O157:H7, the *Z3276* gene (a genetic marker of *E. coli* O157:H7) was detected in only three of these isolates (Fig 1). So, the *E. coli* O157:H7 were identified in 1.1% (3/267) in raw cattle milk.

**FIG 1.**
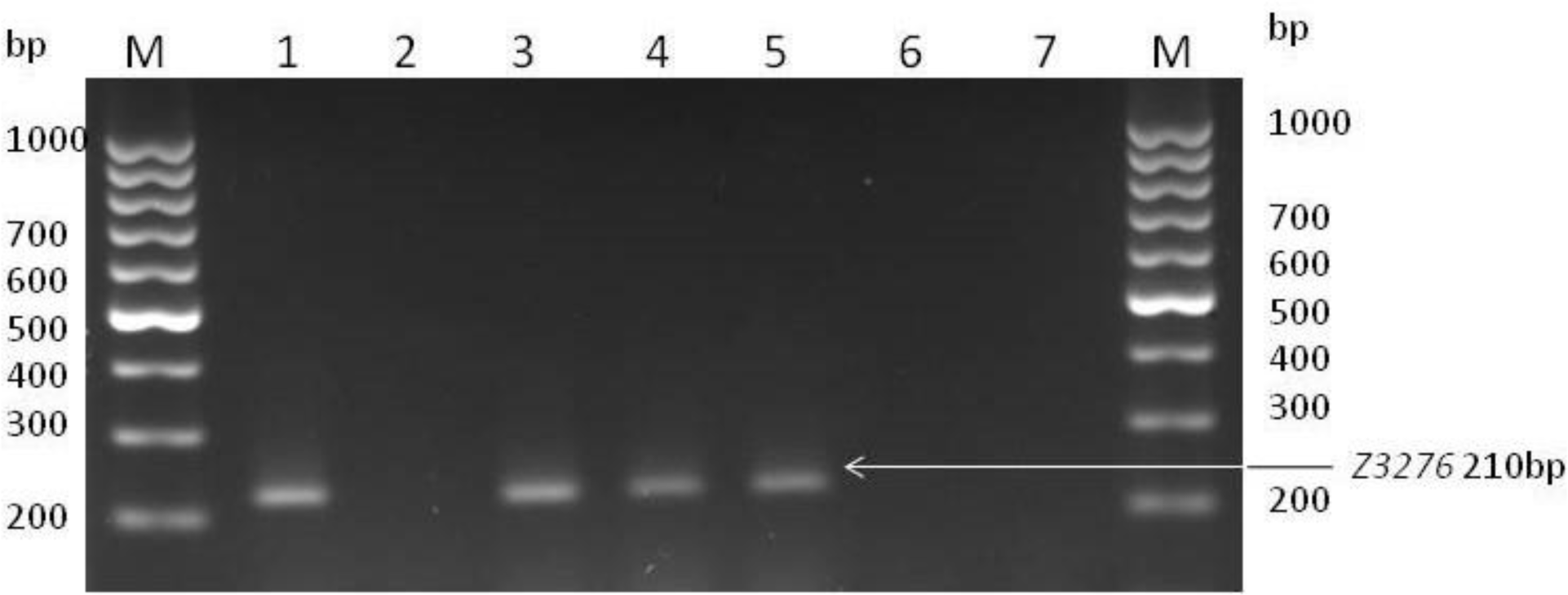
PCR amplication for detection of *Z3276* gene in *E. coli* O157:H7. [Lane M: 100bp ladder, Lane 1: Positive control, Lane 2: Negative control, Lane 3, 4, 5: Positive isolates, Lane 6, 7: Negative isolates].

### Prevalence of O157:H7 STEC and non-O157:H7 STEC

Out of three PCR-confirmed *E. coli* O157:H7, two isolates had both the *stx1* and *stx2* genes and one isolate had only the *stx2* gene (Fig 2). So, the prevalence of O157:H7 STEC is 1.1% (3/267) (Table 1). While the other two biochemically identified O157:H7 isolates, in which the *Z3276* gene was absent, the *stx2* gene was present in both isolates (Fig 2). So, these isolates are sorbitol negative non-O157:H7 STEC (Table 1).

**FIG 2.**
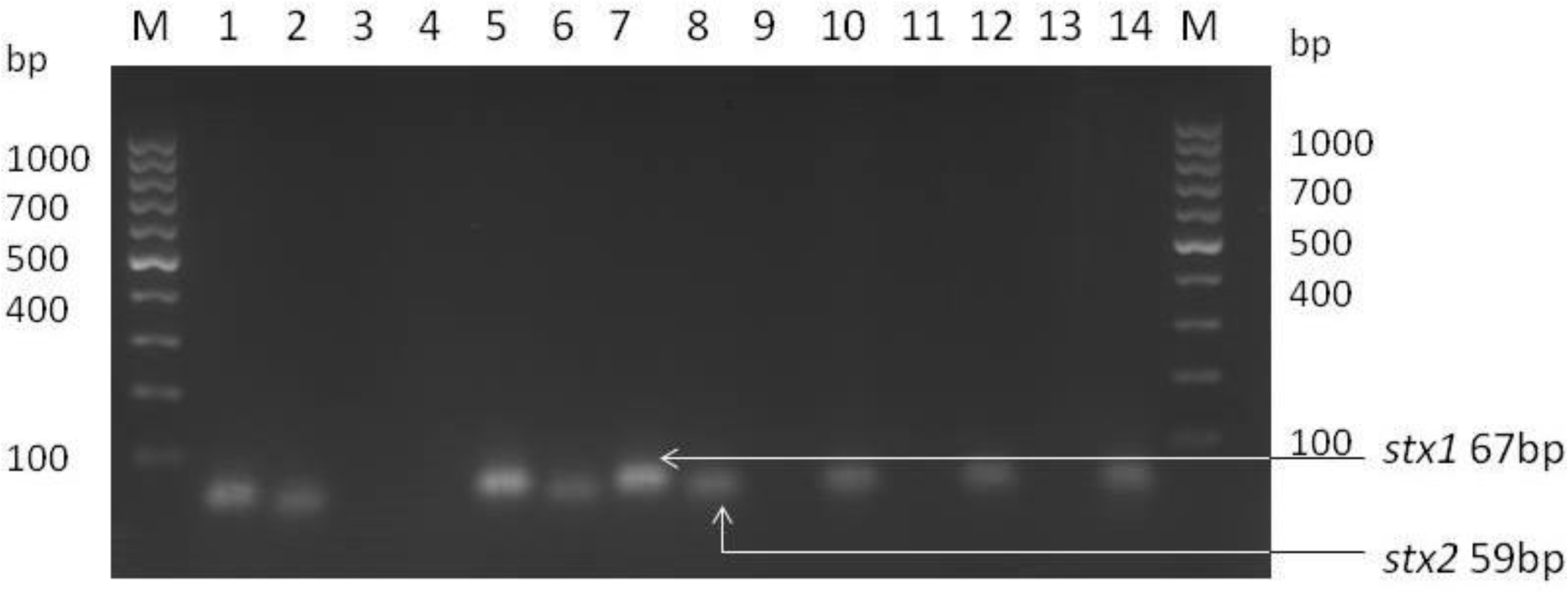
Detection of *stx* genes in *E. coli* O157:H7. [Lane M: 100bp ladder, Lane 1, 2: Positive control, Lane 3, 4: Negative control, Lane 5, 6, 7, 8: *E. coli* O157:H7 with both genes, Lane 9, 10: *E. coli* O157:H7 with only *stx2* gene, Lane 11, 12, 13, 14: Sorbitol negative non-O157:H7 with *stx2* gene].

**FIG 3.**
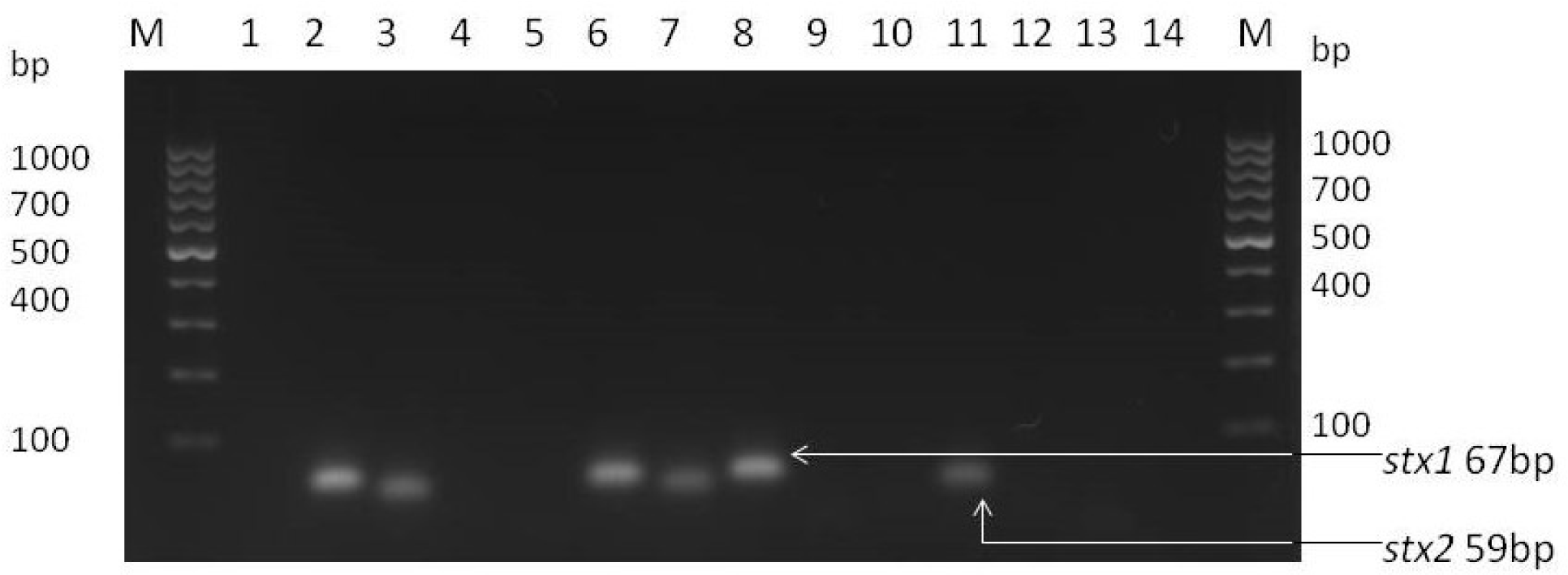
Detection of *stx* genes in non-O157:H7. [Lane M: 100bp ladder, Lane 1, 14: Empty wells, Lane 2, 3: Positive control, Lane 4, 5: Negative control, Lane 6, 7: Isolate with both *stx1* and *stx2* genes, Lane 8, 9: Isolate with only *stx1* gene, Lane 10, 11: Isolate with only *stx2* gene, Lane 12, 13: Negative isolate].

**TABLE 1.** Prevalence of *stx* genes.

| *stx* genes | STEC (number of isolates) |  |  | Total isolates |
| --- | --- | --- | --- | --- |
|  | O157:H7 STEC | Non-O157:H7 STEC | sorbitol negative<br>non-O157:H7 STEC |  |
| *stx1* | 0 | 6 | 0 | 6 |
| *stx2* | 1 | 11 | 2 | 14 |
| *stx1* and *stx2* | 2 | 9 | 0 | 11 |
| Total isolates | 3 | 26 | 2 | 31 |

Out of 74 *E. coli* isolates confirmed biochemically, the *stx1* gene was present in six isolates, the *stx2* was present in 11 isolates and both genes were present in nine isolates. So, the prevalence of non-O157:H7 STEC is 10.5% (26 isolates and two sorbitol-negative non-O157:H7 i.e. 28/267) (Table 1).

### Prevalence of *stx* genes

The overall prevalence of STEC in raw cattle milk is 11.6% (31/267) whereas the prevalence of STEC among *E. coli* isolates is 39.2% (31/79) (Table 1). The prevalence of only *stx1* gene in STEC is 19.3% (6/31; six in non-O157:H7 STEC). The prevalence of only *stx2* gene in STEC is 45.2% (14/31, 11 in non-O157:H7 STEC, one in O157:H7 STEC, and two in sorbitol-negative non-O157:H7 STEC) (Table 1). The prevalence of both genes in total STEC is 35.5% (11/31; nine in non-O157:H7 STEC and two in O157:H7 STEC (Table 1).

### Antibiogram of *E. coli* and STEC

Out of 12 antibiotics tested, 99% (78/79) of *E. coli* isolates were found to be the most sensitive to azithromycin while 61% (48/79) of isolates were found to have an intermediate resistance to ampicillin/sulbactam. Similarly, 18% (14/79) of the *E. coli* isolates showed resistance to tetracycline and doxycycline. A detailed description is provided in Table 2.

**TABLE 2.** Antibiogram profile of *E. coli* and STEC.

| Antibiotics class | Antibiotics | Disc<br>potency<br>(µg) | *E. coli* (n=79) |  |  | STEC (n=31) |  |  |
| --- | --- | --- | --- | --- | --- | --- | --- | --- |
|  |  |  | Sensi<br>tive | Interm<br>ediate | Resis<br>tant | Sensi<br>tive | Interm<br>ediate | Resis<br>tant |
| Aminoglycosides | (Amikacin) AK | 30 | 56<br>(71) | 22 (28) | 1 (1) | 25<br>(81) | 5 (16) | 1 (3) |
|  | (Gentamicin)<br>GEN | 10 | 70<br>(89) | 8 (10) | 1 (1) | 26<br>(84) | 5 (16) | 0 (0) |
|  | (Kanamycin) K | 30 | 36<br>(46) | 43 (54) | 0 (0) | 17<br>(55) | 14 (45) | 0 (0) |
|  | (Streptomycin) S | 10 | 40<br>(51) | 39 (49) | 0 (0) | 18<br>(58) | 13 (42) | 0 (0) |
| Macrolide | (Azithromycin)<br>AZM | 15 | 78<br>(99) | 0 (0) | 1 (1) | 31<br>(100) | 0 (0) | 0 (0) |
| Quinolones | (Enrofloxacin)<br>EX | 5 | 59<br>(75) | 12 (15) | 8 (10) | 28<br>(90) | 1 (3) | 2 (7) |
|  | (Levofloxacin)<br>LE | 5 | 74<br>(94) | 2 (2) | 3 (4) | 31<br>(100) | 0 (0) | 0 (0) |
|  | (Ciprofloxacin)<br>CIP | 5 | 66<br>(84) | 9 (11) | 4 (5) | 26<br>(84) | 5 (16) | 0 (0) |
| Tetracycline | (Tetracycline) TE | 30 | 65<br>(82) | 0 (0) | 14<br>(18) | 27<br>(87) | 0 (0) | 4 (13) |
|  | (Doxycycline)<br>DO | 30 | 65<br>(82) | 0 (0) | 14<br>(18) | 27<br>(87) | 0 (0) | 4 (13) |
| Beta-lactam/beta-lactamase inhibitor | (Ampicillin/<br>Sulbatam) A/S | 10 | 28<br>(35) | 48 (61) | 3 (4) | 9 (29) | 22 (71) | 0 (0) |
| Cephalosporin | (Ceftriaxone)<br>CTR | 30 | 62<br>(79) | 15 (19) | 2 (2) | 23<br>(74) | 6 (19) | 2 (7) |

Note: Figures in parenthesis indicate percentages.

Out of 12 antibiotics tested, 100% (31/31) of STEC isolates were found to be the most sensitive to azithromycin and levofloxacin while 71% (22/31) of isolates were found to have an intermediate response to ampicillin/sulbactam. Similarly, 13% (4/31) of the STEC isolates showed resistance to tetracycline and doxycycline. A detailed description is provided in Table 2.

## DISCUSSION

This study, to the best of our knowledge, revealed for the first time in Nepal that the raw milk in Chitwan District is contaminated with STEC. Thus, raw milk is a potential vector for zoonotic STEC transmission from cattle to humans in this study area resulting in complications like HUS in hospitals. Our findings of the Z3276 gene, *stx1*and *stx2* genes in raw cattle milk in Chitwan clearly indicated a potential risk of STEC to the raw milk consumers in Nepal. The prevalence of *E. coli* reported in this study certainly reveals the overall poor conditions of hygiene/cleanliness of the milk being produced (45) and reveals the chances of faecal contamination as *E. coli* is the best indicator of faecal contamination (46). The finding of zoonotic *E. coli* and STEC is an alarming threat to consumer’s health. The notable higher prevalence of only *stx2* gene established in this study is more concerning as the *stx2* gene is more often linked with the development of Haemorragic Colitis and Hemolytic Uremic Syndrome (30, 47, 48).

We found a higher prevalence of *E. coli* in raw cattle milk in Chitwan, similar to various other studies (23–30%) (49-53). In contrast to our study, much higher prevalence (44–81%) is reported in India, Bangladesh and Ehiopia. This may be due to a difference in collection method from our study causing higher contamination of milk, as they collected milk from a collection tank in Ethiopia (54), a market in Bangladesh (55) and hotels and restaurants in India (56). There are also reports of lower (2–16%) prevalence than our study (57-59). The lower prevalence in other research may be due to different isolation methods, mostly done without enrichment of the sample.

The prevalence of STEC in raw cattle milk in this study is similar to other studies (10– 17%) (49, 60). In this study, the prevalence of STEC O157:H7 was found 1.1%, and non-O157:H7 STEC was found to be 10.5% in raw cattle milk. This finding is similar to other findings in Nigeria and Iran (60, 61). However, studies from China, Pakistan, Bulgaria, and Ethiopia have reported different prevalences of STEC O157:H7 (0–12%) in raw cattle milk (62-65). STEC contamination found in raw milk might be due to sub-clinical mastitis, cross-contamination of milk with faeces, or lack of hygienic measures during the collection and processing of milk (58, 66). The difference in findings may be due to different diagnostic methods used. Tahira used only SMAC while others used latex agglutination test to confirm O157:H7. In other studies, serological confirmation of non-O157:H7 STEC was done. For the rapid and sensitive detection of STEC from clinical and food samples, PCR has proven to be of great diagnostic value (67).

In this study, the prevalence of STEC in *E. coli* isolates was found 39.2% and STEC isolates harbored higher *stx2* genes than *stx1* genes. The finding of our study about even higher isolates with only *stx2* gene than the isolates with both genes suggests that the increased risk of experiencing more severe illness, as the strains that express *stx2* alone are more likely to be associated with the progression to HUS than strains that produce both *stx1* and *stx2* (30). In accordance to our findings, higher *stx2* genes than *stx1* genes are also reported in Iran, Brazil, USA, and Japan (60, 68-70). However, several studies showed *stx1* as the predominant *stx* type (63, 71). The high frequency of *stx2* in faecal samples collected from cows, buffalo, and goats (25, 55) might explain the higher prevalence of *stx2-*positive STEC in raw milk. In humans, epidemiologic data suggest that *E. coli* O157 strains that express *stx2* are more important than *stx1* in the development of HUS (30).

The concerns for the use of antimicrobial products in food-producing animals are increasing as it is directly linked to human food safety because foods of animal origin are sometimes identified as the carriers of resistant food-borne pathogens. The antimicrobial susceptibility testing of *E. coli* and STEC showed that they are mostly sensitive to a majority of the commonly available antibiotics on the market. In this study, *E. coli* and STEC isolates displayed the highest sensitivity to azithromycin. Azithromycin appeared to be useful in decolonizing patients who had recovered from illnesses caused by STEC O104:H4 (72, 73) Similar susceptibility to azithromycin, levofloxacin, gentamicin, and ciprofloxacin was also found in several other studies (74, 75). The results of our study also show varying frequency of antibiotic sensitivity patterns of *E. coli* and STEC in milk as compared to reports of other researchers (22, 65). The prevalence of antibiotic resistance can vary widely based on factors such as antibiotic availability, cost, and the responsible use of these drugs in various parts of the world (76, 77). The differences could also be due to different biochemical and genotypic properties of bacteria, that lead to drug resistance and it spread among different genera and species is mediated by horizontal gene transfer (78, 79).

In this study, the most resistance was seen with tetracycline and doxycycline. Tetracycline is often used as first-line antimicrobial in disease prevention and growth promotion in food animals (80). The high level of resistance to tetracycline may also be due to the widespread use of tetracycline in dairy animals in the treatment of mastitis (81). The *tet A* was the most predominant encoding resistance gene in *E. coli* isolates from cases of clinical and sub-clinical mastitis (82). Similar resistance patterns have been documented in other regions, underlining the global nature of this issue (60, 83, 84). However, *E. coli* did not show any resistance to kanamycin and streptomycin similar to the study in India (85), and in disagreement with the study in Ethiopia (65). Decreased resistance of drugs such as kanamycin and streptomycin in the *E. coli* isolates can be a result of restricted use of these agents in livestock production. The resistant determinants are lost over time without exposure to the drugs (86). This implies selection pressure for *E. coli* in Chitwan for kanamycin resistance has decreased.

This finding of antimicrobial susceptibility suggests that sensitive antibiotics remain effective tools in combating *E. coli* infections in food-producing animals. Although STEC is susceptible to antibiotics, their use should be avoided in its treatment due to the potential risk of hemolytic uremic syndrome. Early detection is essential for the control and prevention of STEC since antibiotics cannot be used for its treatment (36, 38, 39)

## MATERIALS AND METHODS

This cross-sectional study was conducted from December 2018 to October 2019 in Chitwan District, which is located in the south-western part of Bagamati Province of Nepal. The samples were taken from registered dairy co-operatives at different places in Chitwan, Nepal. Multi-stage sampling was done for the collection of samples. First of all, the selection of 10 dairy co-operatives was carried out by simple random sampling. Then the number of households that bring milk to the selected dairy was identified and for every two households one house was selected randomly. From the selected house, one milk sample was collected irrespective of the number of cattle in that house. Approximately 20–30 samples were collected from each dairy. The sample size formula of W. Daniel (87) was used to determine the sample size, which calculated 237 as sample size by considering 0.19 (88) as the expected prevalence. The milk samples were collected according to the protocol of the National Mastitis Council (89) from all teats of dairy cattle. A total of 267 pooled milk samples from all four quarters were collected during milking of cattle. The collected samples were transported to the Veterinary Microbiology Laboratory of Agriculture and Forestry University (AFU), in a cool box with ice and subjected to further processing without storage.

### Isolation and identification of *E. coli*

*E. coli* and *E. coli* O157:H7 were isolated and identified by ISO 16654:2001 (90) standard method with slight modifications. After thawing, subsamples of 3 ml milk from individual 15 ml collected milk sample was kept in 40 ml capacity test tubes containing 27 ml of peptone water (M028, HiMedia, India), homogenized manually, and incubated at 37°C for 24 hrs for non-selective pre-enrichment. For selective enrichment of O157:H7, 1 ml of pre-enriched fluid was taken from the test tubes and added to 9 ml of modified tryptone soya broth (M1286I, HiMedia, India), with novobiocin supplement (FD290, HiMedia, India) and incubated at 41°C for 24 hrs.

For non-O157:H7, following pre-enrichment, a loopful of broth culture was streaked on MacConkey (MAC) agar (M081, HiMedia, India) using three quadrant streak method for a primary culture and incubated at 37°C for 24 hrs. *E. coli* suspected colonies were subcultured in Eosin Methylene Blue (EMB) agar (M317, HiMedia, India) one or more times until a pure colony was obtained. The pure colony was streaked on Nutrient agar (M001, HiMedia, India) in a manner that will allow well-isolated colonies to develop. The pure cultures were subjected to biochemical confirmation.

For O157:H7, following selective enrichment one loopful of broth culture was subcultured onto Sorbitol MacConkey (SMAC) agar (M298I, HiMedia, India) with Cefixime Tellurite supplement (FD147, HiMedia, India). CT-SMAC agar was incubated at 37°C for 24 hrs. Sorbitol-negative colonies from CT-SMAC agar were streaked on EMB agar and typical *E. coli* O157 colonies on the EMB agar were streaked onto Nutrient agar and incubated at 37°C for 24 hrs.

*E. coli* suspected colonies were isolated from raw cattle milk after obtaining the pink colonies on MAC agar that produce green metallic sheen on EMB agar. *E. coli* O157:H7 colonies were identified as pale colonies in CT-SMAC agar as they are sorbitol non-fermenters. The suspected colonies were confirmed to be *E. coli* after performing primary and secondary biochemical tests. The IMViC tests (Indole test, Methyl Red test, Voges Proskauer test, and Citrate utilization test) results have to be positive for Indole, positive for MR test, negative for VP test, and negative for Citrate utilization test. The growth in TSI agar producing acidic slant (yellow), acidic butt (yellow), H_2_S negative, and gas positive were confirmed as *E. coli*. The isolates have to be oxidase and catalase positive. The motility and ornithine decarboxylase activity have to be positive by MIO test.

### Assessment of antibiogram

In vitro antimicrobial sensitivity pattern of identified *E. coli* and STEC was carried out by Kirby-Bauer disc diffusion method (91) on Muller Hinton agar according to the Clinical and Laboratory Standards Institute (92). An inoculum was prepared by incubating five pure colonies in BHI broth (M210, HiMedia, India), then the turbidity was adjusted to 0.5 McFarland standards equivalent to 1.0 × 10^8^ cfu/mL. The inoculum thus prepared was uniformly streaked on Mueller Hinton agar plates using sterile swabs and left for a minute before introduction of the antibiotic discs. Antibiotics (HiMedia, India) commonly used in the field were selected, named as gentamicin (GEN), amikacin (AK), kanamycin (K), streptomycin (S), enrofloxacin (EX), levofloxacin (LE), doxycycline (DO), tetracycline (TE), ciprofloxacin (CIP), ampicillin/ sulbactam (A/S), ceftrizone (CTR) and azithromycin (AZM). The plates were incubated at 37 °C for 24 hrs, and the diameters of the zones of inhibition were measured, and results were interpreted according to CLSI guidelines (92).

### Detection of virulence genes

Isolated *E. coli* strains were investigated for the presence of genetic markers of *E. coli* O157:H7 (*Z3276* gene), and Shiga toxins encoding genes (*stx1* and *stx2* genes). The genomic DNA was extracted from pure cultures of *E. coli* grown overnight in the Nutrient agar at 37°C by using the rapid boiling method (93) with slight modification. The quantification of DNA extracted from *E. coli* was done by measuring absorbance at *A_260_*/*A_280_* ratios using the Spectrophotometer (Quawell, Q5000 UV-Vis Spectrophotometer, USA). The conventional PCR was performed to amplify the genes. The primers used for amplification are described in Table 1. The Thermal Cycler (Bio-Rad T100™, USA) was used for the PCR thermal cycling conditions with a pre-denaturation step at 95°C for 10 min then 40 cycles of denaturation step at 95°C for 10 sec and 40 cycles of annealing temperatures at 60°C for 1 min. After a final elongation step at 60°C for 5 min the sample can be stored at 4°C for infinity (∞). The reaction mixture contained 10 μL platinum green hot start PCR master mix (13001014, Invitrogen, USA), 1 μL of forward primer, 1 μL of reverse primer (FASMAC, Japan) (Table 3), final concentration of 1 μL of DNA template, and 7 μL nuclease free water with the total volume of 20 μL. The amplified PCR products were resolved by electrophoresis tank (Multisub Electrophoresis Systems; Nano PAC - 300P, Cleaver Scientific, USA) in 2% agarose gel (CSL-AG100, Cleaver Scientific, USA) prepared in 1XTBE (Tris-Borate-EDTA) (T4415, Sigma, USA) along with SYBR^TM^ Safe DNA Gel Stain (S33102, Invitrogen, USA). In the first and last wells of the gel, 5 μL of 100bp DNA ladder (SM0243, Thermo Fisher Scientific, USA) was added and 10 μL PCR products were added in the remaining wells along with positive and negative control. The gel electrophoresis was run and set at 90 V, 90 A for 60 min. The gel was taken out from the tray and finally visualized under a UV trans-illuminator (Platinum Q9, Uvitech Cambridge, UK). *E. coli* O157:H7 ATCC 43895 (positive for *stx1* and *stx2* genes) was used as positive control and *Salmonella gallinarum* (SG9R vaccine) was used as negative control.

**TABLE 3.**
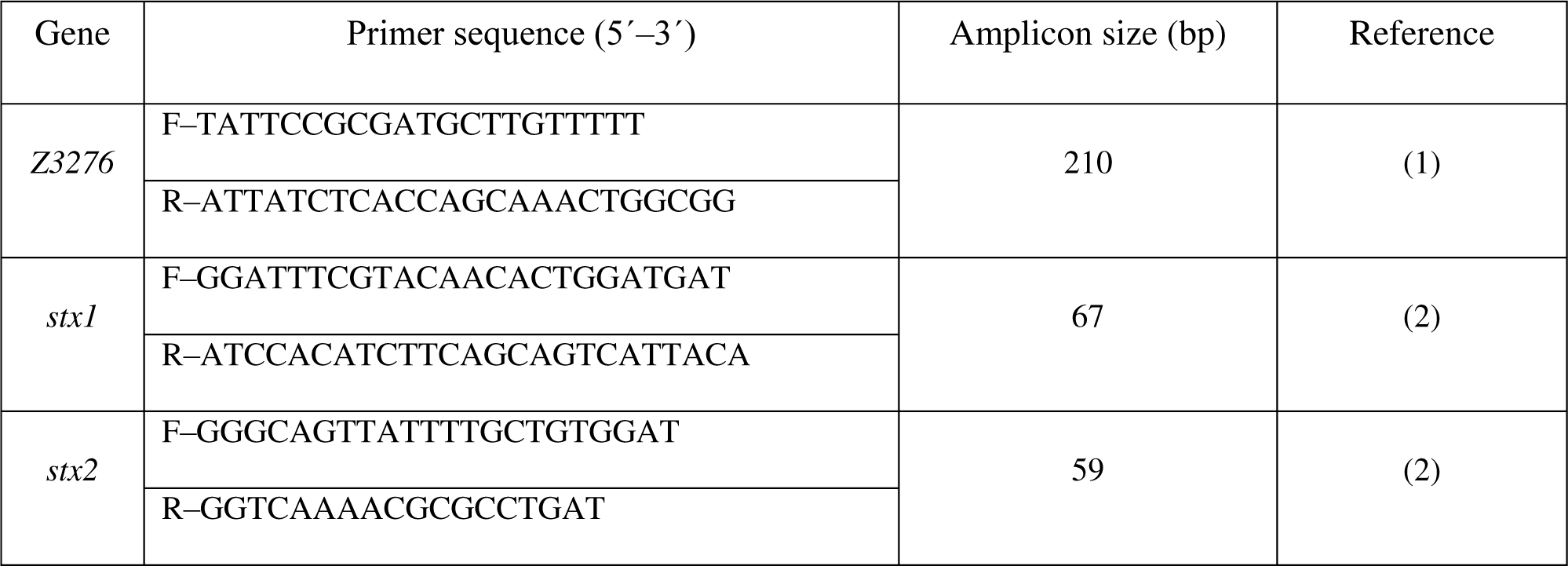
Primers used in this study.

## CONCLUSION

Raw milk is a potential vector for zoonotic STEC transmission from cattle to humans. The occurrence of *E. coli* O157:H7 in milk samples suggests a potential risk of STEC infection due to raw milk consumption in Nepal. The detection of a higher prevalence of *stx2* genes compared to *stx1* genes, or the presence of both genes, shows the risks of greater complications in humans. Our finding of STEC in milk poses a risk for the human population and livestock. As *E. coli* is already established as one of the major causative organisms in sub-clinical mastitis and the higher prevalence of sub-clinical mastitis in Chitwan District makes it more important for molecular-based STEC surveillance in the wider area to minimize the risk of spread of STEC to animal and human. This study on antibiotic susceptibility of *E. coli* and STEC presents a high degree of sensitivity to a range of commonly available antibiotics. This finding suggests that these antibiotics remain effective tools in combating *E. coli* infections in food-producing animals but continuous antimicrobial resistance screening for *E. coli* is necessary to inspect any emerging resistance. When it comes to STEC, despite their sensitivity to antibiotics, they should not be used for treatment due to a lack of evidence supporting their effectiveness and the potential risk of hemolytic uremic syndrome. This makes proper detection of STEC crucial as it cannot be treated with antibiotics, therefore, antibiotics should only be administered in cases after confirming a pathogen that requires such treatment. Nationwide detection of STEC in different sources is essential to implement one health approach of controlling and preventing zoonotic STEC to minimize the risk of health hazards.

## CRediT authorship contribution statement

Sunita Shrestha: Conceptualization, Methodology, Investigation, Writing-original draft. Sirjan Bastola: Conceptualization, Methodology, Resources. Rebanta Kumar Bhattarai: Resources, Validation, Writing-review and editing, Visualization, Supervision. Himal Luitel: Resources, Validation, Supervision. Hom Bahadur Basnet: Resources, Validation, Supervision.

## Acknowledgement

We express our heartfelt gratitude to Adjunct Prof. Dr. Deb Prasad Pandey, Department of Microbiology and Parasitology, FAVF, AFU, for his critical reviews of this paper. Similarly, we are grateful to Dr. Sujata Regmi, Livestock Development Officer, National Livestock Breeding Office, for her help during sample collection, and Biswo Dawadi, laboratory assistant for his help in the molecular study.

## Ethics approval

No ethical permission is required from any committee for the collection of milk samples from cattle according to Nepal law and verbal consent from the farmers was obtained before the collection of milk samples.

## References

1. WHO. 2018. E. coli. https://www.who.int/en/news-room/fact-sheets/detail/e-coli. Accessed 11 June 2019.

2. Bryan A, Youngster I, McAdam AJ. 2015. Shiga toxin producing *Escherichia coli*. Clin Lab Med 35:247–72. 10.1016/j.cll.2015.02.004

3. Mersha G, Asrat D, Zewde BM, Kyule M. 2010. Occurrence of *Escherichia coli* O157:H7 in faeces, skin and carcasses from sheep and goats in Ethiopia. Lett Appl Microbiol 50:71–76. 10.1111/j.1472-765X.2009.02757.x

4. Haileselassie M, Taddele H, Adhana K, Kalayou S. 2013. Food safety knowledge and practices of abattoir and butchery shops and the microbial profile of meat in Mekelle City, Ethiopia. Asian Pac J Trop Biomed 3:407–412. 10.1016/S2221-1691(13)60085-4

5. Signorini M, Tarabla H. 2009. Quantitative risk assessment for verocytotoxigenic *Escherichia coli* in ground beef hamburgers in Argentina. Int J Food Microbiol 132:153–161. 10.1016/j.ijfoodmicro.2009.04.022

6. Fedio WM, Jinneman KC, Yoshitomi KJ, Zapata R, Wendakoon CN, Browning P, Weagant SD. 2011. Detection of *E. coli* O157:H7 in raw ground beef by Pathatrix immunomagnetic-separation, real-time PCR and cultural methods. Int J Food Microbiol 148:87–92. 10.1016/j.ijfoodmicro.2011.05.005

7. Tozzoli R, Scheutz F. 2014. Diarrhoeagenic *Escherichia coli* infections in humans, p 1-18. In Morabito S (ed), Pathogenic Escherichia coli: Molecular and Cellular Microbiology. Caister Academic Press, Rome.

8. Kaper JB, Nataro JP, Mobley HL. 2004. Pathogenic *Escherichia coli*. Nat Rev Microbiol 2:123–140. 10.1038/nrmicro818

9. Mark Taylor C. 2008. Enterohaemorrhagic *Escherichia coli* and *Shigella dysenteriae* type 1-induced haemolytic uraemic syndrome. Pediatr Nephrol 23:1425–1431. 10.1007/s00467-008-0820-3

10. Iweriebor BC, Iwu CJ, Obi LC, Nwodo UU, Okoh AI. 2015. Multiple antibiotic resistances among Shiga toxin producing *Escherichia coli* O157 in feces of dairy cattle farms in Eastern Cape of South Africa. BMC Microbiol 15:1–9. 10.1186/s12866-015-0553-y

11. Motiwala A. 2014. Verotoxigenic *Escherichia coli*: Detection by Commercial Enzyme Immunoassays. *In* Batt CA (ed), Encyclopedia of Food Microbiology. Academic Press, NY, USA.

12. Riley LW, Remis RS, Helgerson SD, McGee HB, Wells JG, Davis BR, Hebert RJ, Olcott ES, Johnson LM, Hargrett NT, Blake PA, Cohen ML. 1983. Hemorrhagic colitis associated with a rare *Escherichia coli* serotype. N Engl J Med 308:681–685. 10.1056/NEJM198303243081203

13. Islam MA. 2009. Shiga toxin-producing *Escherichia coli* in humans and the food chain in Bangladesh. PhD thesis. Wageningen University, Netherlands.

14. Joensen KG, Scheutz F, Lund O, Hasman H, Kaas RS, Nielsen EM, Aarestrup FM. 2014. Real-time whole-genome sequencing for routine typing, surveillance, and outbreak detection of verotoxigenic *Escherichia coli*. J Clin Microbiol 52:1501–1510. 10.1128/JCM.03617-13

15. Perna NT, Plunkett G, 3rd, Burland V, Mau B, Glasner JD, Rose DJ, Mayhew GF, Evans PS, Gregor J, Kirkpatrick HA, Posfai G, Hackett J, Klink S, Boutin A, Shao Y, Miller L, Grotbeck EJ, Davis NW, Lim A, Dimalanta ET, Potamousis KD, Apodaca J, Anantharaman TS, Lin J, Yen G, Schwartz DC, Welch RA, Blattner FR. 2001. Genome sequence of Enterohaemorrhagic *Escherichia coli* O157:H7. Nature 409:529-533. 10.1038/35054089

16. Ravan H, Amandadi M. 2015. Analysis of yeh fimbrial gene cluster in *Escherichia coli* O157:H7 in order to find a genetic marker for this serotype. Curr Microbiol 71:274–82. 10.1007/s00284-015-0842-6

17. Islam MA, Heuvelink AE, de Boer E, Sturm PD, Beumer RR, Zwietering MH, Faruque ASG, Haque R, Sack DA, Talukder KA. 2007. Shiga toxin-producing *Escherichia coli* isolated from patients with diarrhoea in Bangladesh. J Med Microbiol 56:380–385. 10.1099/jmm.0.46916-0

18. CDC. 2022. Reports of selected *E. coli* outbreak investigations https://www.cdc.gov/ecoli/outbreaks.html. Accessed 25 August 2021.

19. Sang WK, Boga HI, Waiyaki PG, Schnabel D, Wamae NC, Kariuki SM. 2012. Prevalence and genetic characteristics of Shigatoxigenic *Escherichia coli* from patients with diarrhoea in Maasailand, Kenya. J Infect Dev Ctries 6:102–108. 10.3855/jidc.1750

20. Neher S, Hazarika AK, Barkalita LM, Borah P, Bora DP, Sharma RK. 2016. Isolation and characterization of Shiga toxigenic *Escherichia coli* of animal and bird origin by multiplex polymerase chain reaction. Vet World 9:123–7. 10.14202/vetworld.2016.123-127

21. Kiranmayi C, Mallika N. 2010. *Escherichia coli* O157:H7 - An emerging pathogen in foods of animal origin. Vet World 3:382. 10.5455/vetworld.2010.382-389

22. Momtaz H, Safarpoor Dehkordi F, Rahimi E, Ezadi H, Arab R. 2013. Incidence of Shiga toxin-producing *Escherichia coli* serogroups in ruminant’s meat. Meat Sci 95:381–388. 10.1016/j.meatsci.2013.04.051

23. Hussein HS, Sakuma T. 2005. Prevalence of Shiga toxin-producing *Escherichia coli* in dairy cattle and their products. J Dairy Sci 88:450–65. 10.3168/jds.s0022-0302(05)72706-5

24. Solomakos N, Govaris A, Angelidis AS, Pournaras S, Burriel AR, Kritas SK, Papageorgiou DK. 2009. Occurrence, virulence genes and antibiotic resistance of *Escherichia coli* O157 isolated from raw bovine, caprine and ovine milk in Greece. Food Microbiol 26:865–871. 10.1016/j.fm.2009.06.002

25. Bandyopadhyay S, Lodh C, Rahaman H, Bhattacharya D, Bera AK, Ahmed FA, Mahanti A, Samanta I, Mondal DK, Bandyopadhyay S, Sarkar S, Dutta TK, Maity S, Paul V, Ghosh MK, Sarkar M, Baruah KK. 2012. Characterization of Shiga toxin producing (STEC) and Enteropathogenic *Escherichia coli* (EPEC) in raw yak (*Poephagus grunniens*) milk and milk products. Res Vet Sci 93:604–610. 10.1016/j.rvsc.2011.12.011

26. Kavyani HR, Shakerian A, Momtaz H, Rahimi E. 2012. The detection of classical enterotoxins of *Staphylococcus aureus* in raw cow milk using the ELISA method. Turkish J Vet Anim Sci 36:319–322. 10.3906/vet-0906-52

27. Popescu A, Angel E. 2009. Analysis of milk quality and its importance for milk processors. Sci P Anim Sci Biotechnol 42:501–506.

28. Thaker H, Brahmbhatt M, Nayak J. 2012. Study on occurrence and antibiogram pattern of *Escherichia coli* from raw milk samples in Anand, Gujarat, India. Vet World 5:556. 10.5455/vetworld.2012.556-559

29. Mungai EA, Behravesh CB, Gould LH. 2015. Increased outbreaks associated with nonpasteurized milk, United States, 2007-2012. Emerg Infect Dis 21:119-22. 10.3201/eid2101.140447

30. Farrokh C, Jordan K, Auvray F, Glass K, Oppegaard H, Raynaud S, Thevenot D, Condron R, De Reu K, Govaris A, Heggum K, Heyndrickx M, Hummerjohann J, Lindsay D, Miszczycha S, Moussiegt S, Verstraete K, Cerf O. 2013. Review of Shiga-toxin-producing *Escherichia coli* (STEC) and their significance in dairy production. Int J Food Microbiol 162:190–212. 10.1016/j.ijfoodmicro.2012.08.008

31. Magwira CA, Gashe BA, Collison EK. 2005. Prevalence and antibiotic resistance profiles of *Escherichia coli* O157:H7 in beef products from retail outlets in Gaborone, Botswana. J Food Prot 68:403–406. 10.4315/0362-028x-68.2.403

32. Rasheed MU, Thajuddin N, Ahamed P, Teklemariam Z, Jamil K. 2014. Antimicrobial drug resistance in strains of *Escherichia coli* isolated from food sources. Rev Inst Med Trop Sao Paulo 56:341–346. 10.1590/s0036-46652014000400012

33. Effendi MH, Harijani N. 2017. Profile antibiotics resistance on *Escherichia coli* isolated from raw milk In Surabaya Dairy Farms, Indonesia. TOJDAC 7:1340–1344. 10.7456/1070DSE/108

34. Bruyand M, Mariani-Kurkdjian P, Gouali M, de Valk H, King LA, Le Hello S, Bonacorsi S, Loirat C. 2018. Hemolytic uremic syndrome due to Shiga toxin-producing *Escherichia coli* infection. Med Mal Infect 48:167–174. 10.1016/j.medmal.2017.09.012

35. Muhlen S, Dersch P. 2020. Treatment strategies for infections with Shiga toxin-producing *Escherichia coli*. Front Cell Infect Microbiol 10:169. 10.3389/fcimb.2020.00169

36. Wong CS, Jelacic S, Habeeb RL, Watkins SL, Tarr PI. 2000. The risk of the hemolytic-uremic syndrome after antibiotic treatment of *Escherichia coli* O157:H7 infections. N Engl J Med 342:1930–6. 10.1056/NEJM200006293422601

37. Zhang X, McDaniel AD, Wolf LE, Keusch GT, Waldor MK, Acheson DW. 2000. Quinolone antibiotics induce Shiga toxin-encoding bacteriophages, toxin production, and death in mice. J Infect Dis 181:664–670. 10.1086/315239

38. Freedman SB, Xie J, Neufeld MS, Hamilton WL, Hartling L, Tarr PI, Alberta Provincial Pediatric Enteric Infection T, Nettel-Aguirre A, Chuck A, Lee B, Johnson D, Currie G, Talbot J, Jiang J, Dickinson J, Kellner J, MacDonald J, Svenson L, Chui L, Louie M, Lavoie M, Eltorki M, Vanderkooi O, Tellier R, Ali S, Drews S, Graham T, Pang XL. 2016. Shiga toxin-producing *Escherichia coli* infection, antibiotics, and risk of developing Hemolytic Uremic Syndrome: A Meta-analysis. Clin Infect Dis 62:1251-1258. 10.1093/cid/ciw099

39. Tarr PI, Freedman SB. 2022. Why antibiotics should not be used to treat Shiga toxin-producing *Escherichia coli* infections. Curr Opin Gastroenterol 38:30–38. 10.1097/MOG.0000000000000798

40. Schroeder CM, Zhao C, DebRoy C, Torcolini J, Zhao S, White DG, Wagner DD, McDermott PF, Walker RD, Meng J. 2002. Antimicrobial resistance of *Escherichia coli* O157 isolated from humans, cattle, swine, and food. Appl Environ Microbiol 68:576–581. 10.1128/AEM.68.2.576-581.2002

41. MoAD. 2015/16. Statistical Information on Nepalese Agriculture,. https://moald.gov.np/wp-content/uploads/2022/04/STATISTICAL-INFORMATION-ON-NEPALESE-AGRICULTURE-2072-73.pdf. Accessed 7 March 2018.

42. RSS. 2019. Chitwan becomes self-sufficient in milk, p *In* myRepublica. https://myrepublica.nagariknetwork.com/tag/Veterinary_Hospital.

43. Bhandari S, Subedi D, Tiwari BB, Shrestha P, Shah S, Al-Mustapha AI. 2021. Prevalence and risk factors for multidrug-resistant *Escherichia coli* isolated from sub-clinical mastitis in the western Chitwan region of Nepal. J Dairy Sci 104:12765–12772. 10.3168/jds.2020-19480

44. Pandit A, Khanal T. 2013. Assessment of sub-clinical mastitis and its associated risk factors in dairy livestock of Lamjung, Nepal. Int J Infect Microbiol 2:49–54. 10.3126/ijim.v2i2.8322

45. Meshref AMS. 2013. Bacteriological quality and safety of raw cow’s milk and fresh cream. Slov Vet Res 50:21–30.

46. National Research Council. 1985. An evaluation of the role of microbiological criteria for foods and food ingredients. The National Academies Press, Washington, DC.

47. Etcheverria AI, Padola NL. 2013. Shiga toxin-producing *Escherichia coli*: factors involved in virulence and cattle colonization. Virulence 4:366–72. 10.4161/viru.24642

48. Manning SD, Motiwala AS, Springman AC, Qi W, Lacher DW, Ouellette LM, Mladonicky JM, Somsel P, Rudrik JT, Dietrich SE, Zhang W, Swaminathan B, Alland D, Whittam TS. 2008. Variation in virulence among clades of *Escherichia coli* O157:H7 associated with disease outbreaks. Proc Natl Acad Sci U S A 105:4868–73. 10.1073/pnas.0710834105

49. Ranjbar R, Safarpoor Dehkordi F, Sakhaei Shahreza MH, Rahimi E. 2018. Prevalence, identification of virulence factors, O-serogroups and antibiotic resistance properties of Shiga-toxin producing *Escherichia coli* strains isolated from raw milk and traditional dairy products. Antimicrob Resist Infect Control 7:53. 10.1186/s13756-018-0345-x

50. Bedasa S, Shiferaw D, Abraha A, Moges T. 2018. RETRACTED ARTICLE: Occurrence and antimicrobial susceptibility profile of *Escherichia coli* O157:H7 from food of animal origin in Bishoftu town, Central Ethiopia. Int JFood Contam 5:1–8. 10.1186/s40550-018-0064-3

51. Sheikh JA, Rashid M, Rehman MU, Bhat MA. 2013. Occurrence of multidrug resistance shiga-toxin producing *Escherichia coli* from milk and milk products. Vet World 6:915–918. 10.14202/vetworld.2013.915-918

52. Farhan R, Abdalla S, Abdelrahaman HA, Fahmy N, Salama E. 2014. Prevalence of *Escherichia coli* in some selected foods and children ctools with special reference to molecular characterization of enterohemorrhagic strain. Am J Anim Vet Sci 9:245–251. 10.3844/ajavsp.2014.245.251

53. Elbagory A-RM, Hammad AM, Shiha AMA. 2015. Prevalence of coliforms, antibiotic resistant coliforms and *E. coli* serotypes in some varieties of raw milk cheese in Egypt. Nutr Food Technol. 10.16966/2470-6086.114

54. Shunda D, Habtamu T, Endale B. 2013. Assessment of bacteriological quality of raw cow milk at different critical points in Mekelle, Ethiopia. Int J Livest Res 3:42–48.

55. Islam MA, Mondol AS, de Boer E, Beumer RR, Zwietering MH, Talukder KA, Heuvelink AE. 2008. Prevalence and genetic characterization of Shiga toxin-producing *Escherichia coli* isolates from slaughtered animals in Bangladesh. Appl Environ Microbiol 74:5414–21. 10.1128/AEM.00854-08

56. Bhoomika, Shakya S, Patyal A, Gade NE. 2016. Occurrence and characteristics of extended-spectrum beta-lactamases producing *Escherichia coli* in foods of animal origin and human clinical samples in Chhattisgarh, India. Vet World 9:996–1000. 10.14202/vetworld.2016.996-1000

57. Zeinhom M. 2011. Monitoring of enteric pathogens in milk and some dairy products with special reference for *Enterobacter sakazakii* and *E. coli* O157: H7. PhD thesis. Beni-Suef University, Egypt.

58. Elmonir W, Abo-Remela E, Sobeih A. 2018. Public health risks of *Escherichia coli* and *Staphylococcus aureus* in raw bovine milk sold in informal markets in Egypt. J Infect Dev Ctries 12:533–541. 10.3855/jidc.9509

59. Martha MS, Adelard BM, Lughano JK, Neema K. 2016. Prevalence and antibiotic susceptibility of *Escherichia coli* and *Salmonella* spp. isolated from milk of zero grazed cows in Arusha city. Afr J Microbiol Res 10:1944–1951.

60. Mohammadi P, Abiri R, Rezaei M, Salmanzadeh-Ahrabi S. 2013. Isolation of Shiga toxin-producing *Escherichia coli* from raw milk in Kermanshah, Iran. Iran J Microbiol 5:233–238.

61. Yakubu Y, Shuaibu AB, Ibrahim AM, Hassan UL, Nwachukwu RJ. 2018. Risk of Shiga toxigenic *Escherichia coli* O157:H7 infection from raw and fermented milk in Sokoto Metropolis, Nigeria. J Pathog 2018. 10.1155/2018/8938597

62. Chao G, Zhou X, Jiao X, Qian X, Xu L. 2007. Prevalence and antimicrobial resistance of foodborne pathogens isolated from food products in China. Foodborne Pathog Dis 4:277–284. 10.1089/fpd.2007.0088

63. Tahira B, Ullah K, Samad A, Ali S, Nabi S, Naeem M, Hussain H, Mohammad A. 2017. Isolation and molecular characterization of Shiga toxin producing *E. coli* O157: H7 in raw milk using mPCR. Int J Pharm Sci Res 8:3107–3112. 10.13040/IJPSR.0975-8232.8

64. Koev K, Zhelev G, Marutsov P, Gospodinova K, Petrov V. 2018. Isolation and primary identification of Shiga toxin producing *Escherichia coli* O157 in dairy cattle. Bulg J Vet Med 21:445–450. 10.15547/bjvm.1077

65. Disassa N, Sibhat B, Mengistu S, Muktar Y, Belina D. 2017. Prevalence and antimicrobial susceptibility pattern of *E. coli* O157:H7 isolated from traditionally marketed raw cow milk in and around Asosa Town, Western Ethiopia. Vet Med Int 2017:7581531. 10.1155/2017/7581531

66. Ahmadi E, Mardani K, Amiri A. 2020. Molecular detection and antimicrobial resistance patterns of Shiga toxigenic *Escherichia coli* isolated from bovine sub-clinical mastitis milk samples in Kurdistan, Iran. Arch Razi Inst 75:169–177. 10.22092/ari.2019.124238.1278

67. Paton AW, Paton JC. 2005. Multiplex PCR for direct detection of Shiga toxigenic *Escherichia coli* strains producing the novel subtilase cytotoxin. J Clin Microbiol 43:2944–2947. 10.1128/JCM.43.6.2944-2947.2005

68. Vendramin T, Kich DM, Molina RD, Souza CFVd, Salvatori RU, Pozzobon A, Bustamante-filho IC. 2014. Molecular screening of bovine raw milk for the presence of Shiga toxin-producing *Escherichia coli* (STEC) on dairy farms. Food Sci Technol 34:604–608. 10.1590/1678-457x.6422

69. Zhao T, Doyle MP, Shere J, Garber L. 1995. Prevalence of Enterohemorrhagic *Escherichia coli* O157:H7 in a survey of dairy herds. Appl Environ Microbiol 61:1290–1293. 10.1128/aem.61.4.1290-1293.1995

70. Keen JE, Elder RO. 2002. Isolation of Shiga-toxigenic *Escherichia coli* O157 from hide surfaces and the oral cavity of finished beef feedlot cattle. J Am Vet Med Assoc 220:756–763. 10.2460/javma.2002.220.756

71. Chiueh LC, Liu FM, Shih DYC. 2020. Prevalence of Shiga toxin-producing *Escherichia coli* in feces and raw milk of domestic cattle and sheep. J Food Drug Anal 10:39–46. 10.38212/2224-6614.2769

72. CFSPH. 2016. Enterohemorrhagic *Escherichia coli* and other *E. coli* causing Hemolytic Uremic Syndrome. https://www.cfsph.iastate.edu/Factsheets/pdfs/e_coli.pdf. Accessed 19 February 2019.

73. Nitschke M, Sayk F, Hartel C, Roseland RT, Hauswaldt S, Steinhoff J, Fellermann K, Derad I, Wellhoner P, Buning J, Tiemer B, Katalinic A, Rupp J, Lehnert H, Solbach W, Knobloch JK. 2012. Association between azithromycin therapy and duration of bacterial shedding among patients with Shiga toxin-producing Enteroaggregative *Escherichia coli* O104:H4. JAMA 307:1046–52. 10.1001/jama.2012.264

74. Islam MA, Mondol AS, Azmi IJ, de Boer E, Beumer RR, Zwietering MH, Heuvelink AE, Talukder KA. 2010. Occurrence and characterization of Shiga toxin-producing *Escherichia coli* in raw meat, raw milk, and street vended juices in Bangladesh. Foodborne Pathog Dis 7:1381–5. 10.1089/fpd.2010.0569

75. Tanzin T, Nazir K, Zahan M, Parvej M, Zesmin K, Rahman M. 2016. Antibiotic resistance profile of bacteria isolated from raw milk samples of cattle and buffaloes. J Adv Vet Anim Res 3:62. 10.5455/javar.2016.c133

76. Ghimire K, Banjara MR, Marasini BP, Gyanwali P, Poudel S, Khatri E, Dhimal M. 2023. Antibiotics prescription, dispensing practices and antibiotic resistance pattern in common pathogens in Nepal: A Narrative Review. Microbiol Insights 16:11786361231167239. 10.1177/11786361231167239

77. Palma E, Tilocca B, Roncada P. 2020. Antimicrobial resistance in veterinary medicine: An Overview. Int J Mol Sci 21:1914. 10.3390/ijms21061914

78. Bentley R, Meganathan R. 1982. Biosynthesis of vitamin K (menaquinone) in bacteria. Microbiol Rev 46:241–280. 10.1128/mr.46.3.241-280.1982

79. Schwarz S, Kehrenberg C, Walsh TR. 2001. Use of antimicrobial agents in veterinary medicine and food animal production. Int J Antimicrob Agents 17:431–437. 10.1016/s0924-8579(01)00297-7

80. Granados-Chinchilla F, Rodriguez C. 2017. Tetracyclines in Food and Feedingstuffs: From Regulation to Analytical Methods, Bacterial Resistance, and Environmental and Health Implications. J Anal Methods Chem 2017:1315497. 10.1155/2017/1315497

81. El-Razik KAA, Arafa AA, Hedia RH, Ibrahim ES. 2017. Tetracycline resistance phenotypes and genotypes of coagulase-negative staphylococcal isolates from bubaline mastitis in Egypt. Vet World 10:702–710. 10.14202/vetworld.2017.702-710

82. Messele YE, Abdi RD, Tegegne DT, Bora SK, Babura MD, Emeru BA, Werid GM. 2019. Analysis of milk-derived isolates of *E. coli* indicating drug resistance in central Ethiopia. Trop Anim Health Prod 51:661–667. 10.1007/s11250-018-1737-x

83. Rangel P, Marin JM. 2009. Analysis of *Escherichia coli* isolated from bovine mastitic milk. Pesq Vet Bras 29:363–368. 10.1590/s0100-736x2009000500001

84. Tark DS, Moon DC, Kang HY, Kim SR, Nam HM, Lee HS, Jung SC, Lim SK. 2017. Antimicrobial susceptibility and characterization of extended-spectrum beta-lactamases in *Escherichia coli* isolated from bovine mastitic milk in South Korea from 2012 to 2015. J Dairy Sci 100:3463–3469. 10.3168/jds.2016-12276

85. Sharma S, Khan A, Dahiya DK, Jain J, Sharma V. 2015. Prevalence, identification and drug resistance pattern of *Staphylococcus aureus* and *Escherichia coli* isolated from raw milk samples of Jaipur city of Rajasthan. J Pure Appl Microbiol 9:341–348.

86. Giguere S, Prescott JF, Dowling PM. 2013. Antimicrobial therapy in veterinary medicine, 5^th^ ed. John Wiley & Sons, Hoboken, NJ, USA.

87. Daniel W. 1999. Biostatistics: A Foundation for analysis in the health sciences, 7^th^ ed, vol 141. John Wiley & Sons, Hoboken, NJ, USA.

88. Sethulekshmi C, Latha C, Anu CJ. 2018. Occurrence and quantification of Shiga toxin-producing *Escherichia coli* from food matrices. Vet World 11:104–111. 10.14202/vetworld.2018.104-111

89. Middleton JR, Saeman A, Fox LK, Lombard J, Hogan JS, Smith KL. 2014. The National Mastitis Council: A Global Organization for Mastitis Control and Milk Quality, 50 Years and Beyond. J Mammary Gland Biol Neoplasia 19:241-51. 10.1007/s10911-014-9328-6

90. ISO. 2001. Microbiology of food and animal feeding stuffs - Horizontal method for the detection of *Escherichia coli* O157. ISO 16654:2001, Geneva, Switzerland.

91. Bauer AW, Kirby WMM, Sherris JC, Turck M. 1966. Antibiotic susceptibility testing by a standardized single disk method. Am J Clin Pathol 45:493–496. 10.1093/ajcp/45.4_ts.493

92. CLSI. 2018. Performance Standards for Antimicrobial Disk Susceptibility Tests; Approved Standard, 12^th^ ed. Clinical and Laboratory Standards Institute.

93. Diana J, Pui C, Son R. 2012. Enumeration of *Salmonella* spp., Salmonella typhi and Salmonella typhimurium in fruit juices. Int Food Res J 19:51–56.

## References

1. Li B, Hu Z, Elkins CA. 2014. Detection of live *Escherichia coli* O157:H7 cells by PMA-qPCR. J Vis Exp 84:e50967. 10.3791/50967

2. Li B, Liu H, Wang W. 2017. Multiplex real-time PCR assay for detection of *Escherichia coli* O157:H7 and screening for non-O157 Shiga toxin-producing *E. coli*. BMC Microbiol 17:215. 10.1186/s12866-017-1123-2

